# C19ORF66 broadly escapes viral-induced endonuclease cleavage and restricts Kaposi’s Sarcoma Associated Herpesvirus (KSHV)

**DOI:** 10.1101/506410

**Authors:** William Rodriguez, Aman Srivastav, Mandy Muller

## Abstract

One striking characteristic of certain herpesviruses is their ability to induce rapid and widespread RNA decay in order to gain access to host resources. This phenotype is induced by viral endoribonucleases, including SOX in KSHV, muSOX in MHV68, BGLF5 in EBV and vhs in HSV-1. Here, we performed comparative RNA-seq upon expression of these herpesviral endonucleases in order to characterize their effect on the host transcriptome. Consistent with previous reports, we found that approximately two thirds of transcripts are downregulated in cells expressing any of these viral endonucleases. Among transcripts spared from degradation, we uncovered a cluster of transcripts that systematically escape degradation from all tested endonucleases. Among these escapees, we identified C19ORF66 and reveal that like the previously identified escapees, this transcript is protected from degradation by its 3’UTR. We then show that C19ORF66, a known anti-viral protein, is a potent KSHV restriction factor, suggesting that its ability to escape viral cleavage may be an important component of the host response to viral infection. Collectively, our comparative approach is a powerful tool to pinpoint key regulators of the viral-host interplay and led us to uncover a novel KSHV regulator.

## Introduction

Many viruses including alpha- and gammaherpesviruses, influenza A virus, and SARS coronavirus induce widespread mRNA decay through the use of virally encoded endonucleases (1-5). This process, known as “host shutoff”, allows viruses to rapidly restrict gene expression in order to dampen immune responses and provide access to the host’s resources for viral replication (2,6,7). One well-studied viral endonuclease is the SOX protein encoded by Kaposi’s sarcoma-associated herpesvirus (KSHV). SOX is conserved throughout the herpesvirus family, but only gammaherpesviral SOX homologs display ribonuclease activity in cells (8-10) and studies indicate that SOX activity is important for the *in-vivo* viral lifecycle (11,12). Although SOX targets a degenerate RNA motif present on most mRNA (13-15), multiple studies have shown that some transcripts robustly escape SOX-induced decay (16-21). Studying these ‘escapees’ in aggregate is complicated, however, by the fact that multiple mechanisms can promote apparent escape. These include lack of a targeting motif, indirect transcriptional effects, and active evasion of ribonucleolytic cleavage (16,20-24). This latter phenotype, termed “dominant escape”, is particularly notable as it involves a specific RNA element whose presence in the 3’ UTR of an mRNA protects against SOX cleavage, regardless of whether the RNA contains a targeting motif (19-21). This protective RNA element was termed SRE (for <u>S</u>OX <u>R</u>esistance <u>E</u>lement), but we recently showed that the SRE is also effective against a broad range of viral endonucleases. Perhaps more surprisingly, the SRE is unable to restrict endonucleolytic cleavage originating from a cellular endonuclease, making it the first identified viral-specific ribonuclease escape element(19). We showed that this broad-acting RNA element is not characterized by a defined sequence motif (19) rendering it difficult to identify new escaping transcripts by traditional sequence search. Consequently, the host vs. viral endonuclease dichotomy to only defining characteristic of this novel type of RNA element.

Little is currently known about these types of RNA elements; how widespread they may be in the genome and how they may contribute to the overall viral-host arms race for the control of resources. To date, only two SRE-bearing dominant escapees are known: the host interleukin-6 (IL-6) (18,20,21) and the growth arrest and DNA damage-inducible 45 beta (GADD45B) (19) transcripts. Both the IL-6 and GADD45B SREs were mapped to their 3’UTR and were shown to protect against an array of viral – but not host – RNAses. Furthermore, while little sequence homology was detected among these SREs, we showed that they share similarity in their secondary structure; reinforcing the idea the SRE may function as a platform to recruit a protective protein complex as previously observed (19-21). Functionally, while the beneficial role of IL-6 for KSHV during infection is well documented (25-33), the role of GADD45B is still unclear. In fact, GADD45B is repressed during KSHV latency (34) and GADD45B known pro-apoptotic roles may indicate that this transcript escapes to participate in an anti-viral response to host shutoff.

Here, taking advantage of the ability of the SRE element to block decay from a diverse set of viral endonucleases, we sought to identify novel escaping mRNAs containing SRE or SRE-like elements in the transcriptome. Using comparative RNA-seq, we uncovered a cluster of 75 host mRNAs that escape degradation from four herpesviral endonucleases. Similarly to the previously identified SRE-bearing transcripts, these transcripts were spared from a range of viral – but not host – endonucleases, further supporting that our approach successfully identified novel dominant escapees. Among this list of newly identified escapees, we demonstrate that our top candidate, C19ORF66, is a negative regulator of the KSHV life cycle.

C19orf66 (also annotated RyDEN, IRAV, and SVA-1) is an interferon stimulated gene (ISG) that has been found to be upregulated upon infection by a number a viruses (35-40), including herpesviruses (41,42) in several large scale screens. Recently, C19ORF66 was demonstrated to repress Dengue Virus (DENV) replication and gene expression by interacting with the cytoplasmic poly-A binding protein, PABPC (43), and the RNA helicase MOV10 (44) suggesting that C19ORF66 may restrict DENV infection by either directly influencing the host gene expression machinery, and/or directly targeting viral RNA for degradation, making it an intriguing candidate dominant escapees during KSHV infection.

Here we show that C19ORF66 is upregulated during KSHV infection and accumulates over the course of 96 hours post-reactivation. Knocking down C19ORF66 during KSHV infection leads to higher expression levels of early and delayed early viral genes, which results in higher yields of infectious viral particles and suggests that C19ORF66 has anti-viral activity on KSHV. Taken together, these results demonstrate that SRE and SRE-like elements may be more common than anticipated in the genome, and that transcripts encoding these escape elements may also function as viral restriction factors.

## Results

### Comparative RNA-seq identifies a cluster of common escaping transcripts

Prior analyses indicated that certain host mRNA transcripts robustly escape viral-induced RNA decay by encoding an RNA element in their 3’UTRs. We demonstrated that this RNA element, herein referred to as SRE (**S**OX **R**esistance **E**lement), provides protection against KSHV SOX as well as a variety of viral endonucleases. To identify mRNA transcripts containing SRE or SRE-like elements, we performed comparative RNA-seq based transcriptomics analyses upon expression of the herpesviral RNA endonucleases. Pure populations of cells expressing either KSHV SOX, MHV68 muSOX, Epstein-Barr Virus (EBV) BGLF5, Herpes Simplex 1 (HSV-1) vhs or an empty vector control were generated using Thy1.1-based cell sorting as described before (45). Total RNA was extracted, polyA enriched and cDNA were generated. cDNA libraries were sequenced with a 100-base single-end read on an Illumina HiSeq4000. Resulting reads were aligned to the human genome (hg38) using Bowtie, replicates were merged using CuffCompare and significant expression fold change between mock and each of the endonuclease conditions were assessed by CuffDiff (**Figure 1, Figure S1 & Table S1**). The reproducibility between replicate experiments was high (**Fig. 1A-D**), which is in line with previous reports showing that these endonucleases target transcripts in a selective/sequence-specific manner as previously observed (13). As expected, a number of transcripts were significantly affected upon expression of the various herpesviral endonucleases (**Fig. S1**): we observed that between 55-60% of total mRNA were degraded, with muSOX being the most effective of the endonucleases tested here (**Fig. S1**). This rate of degradation is within range of what was observed before (16). Gene Ontology (GO) analysis on the transcripts spared from degradation revealed that they encode proteins that have a wide array of functions, ranging from ion binding to RNA binding (**Fig S2**).

**Figure 1:**
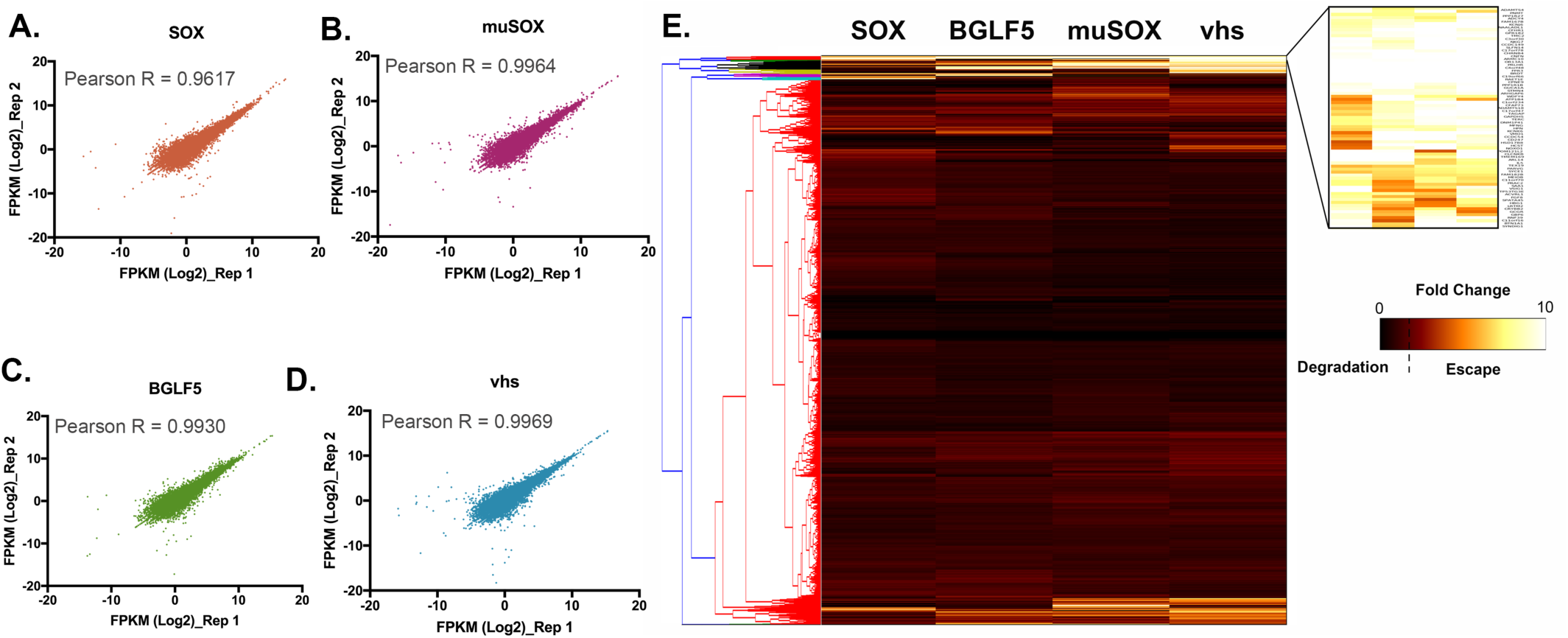
Comparative RNA-seq of the herpesviral RNA endonuclease. **(A-D)** Scatter plots to compare gene expression expressed as log 2 FPKM (Fragments Per Kilobase of transcript per Million mapped reads) amongst replicate experiments. The Pearson correlation coefficient, R, is shown for each plot. **(E)** Hierarchical clutering and heatmap of RNA-seq data: Fold change in expression levels for each condition (SOX, muSOX, BGLF5 and vhs – columns) over mock were normalized are represented as a heatmap. Transcripts were clustered by similarity using the complete linkage method (dendogram on the left). A cluster representing transcript escaping degradation by all tested endonucleases emerged and is enlarged on the top right corner.

To identify transcripts that escape degradation from all 4 endonucleases, we performed hierarchical clustering on the transcript expression data. **Figure 1E** shows a heatmap of the correlation matrix across all transcripts. A cluster encompassing 75 transcripts (**Table S2**) represents the mRNA that escape degradation from all 4 herpesviral endonucleases. We hypothesize that this cluster of transcripts is likely to include mRNA containing SRE or SRE-like elements.

### Candidate escapees are broadly protected from cleavage by viral but not cellular endonucleases

We next set out to investigate further this cluster of common escapees. The RNA-seq hits identified by hierarchical clustering were ranked by confidence (reproducibility among experimental replicates and escaped all endonucleases in all replicates). To confirm the RNA-seq data, we first examined whether the top 10% (**Table S2**) of this list of common escapees were resistant to host shutoff upon lytic reactivation of a KSHV-positive renal carcinoma cell line stably expressing the KSHV BAC16 (iSLK.219). iSLK.219 cells harbor a doxycycline (dox)-inducible version of the major viral lytic transactivator RTA which promotes entry into the lytic cycle upon doxycycline treatment (46,47). As opposed to the housekeeping gene GAPDH that is naturally susceptible to host shutoff, we observed that the mRNA levels of these candidate SRE-bearing mRNAs remained unchanged in reactivated iSLK.219 cells as measured by RT-qPCR (**Fig. 2A**) confirming that these transcripts are resistant to host shutoff in lytically infected cells. Additionally, we recently showed that SRE-containing transcripts are resistant to endonucleases beyond the herpesvirus family (19). We next tested the ability of these novel escapees to evade the heterologous host shutoff from the influenza A virus endonuclease (IAV; PA-X). As shown in **Figure 2B**, contrary to GAPDH, the candidate transcripts were resistant to all endonucleases tested, including PA-X. Finally, one characteristic of SRE-containing mRNAs is that they are still susceptible to cleavage by cellular endonucleases (19). To test whether this was also the case for our novel candidate SRE-bearing transcripts, we monitored cleavage upon expression of the nsp1 protein from SARS coronavirus. Nsp1 is not a nuclease but rather activates mRNA cleavage by an as yet unknown cellular endonuclease *via* a mechanism reminiscent of no-go decay (48,49). Nsp1 thus allows us to induce RNA decay using a viral trigger but carried out by a cellular endonuclease. Nsp1 was transfected into 293T cells and depletion of the candidate transcripts was measured by RT-qPCR. Similar to what we observed before, the candidate escapee mRNAs were not protected in nsp1-expressing cells (**Fig. 2C**). Collectively, these results suggest that the escaping mRNAs identified in our comparative RNA-seq dataset are broadly protected against viral but not cellular endonucleases and we predict that these transcripts may contain an SRE or an SRE-like element that provides broad protection.

**Figure 2:**
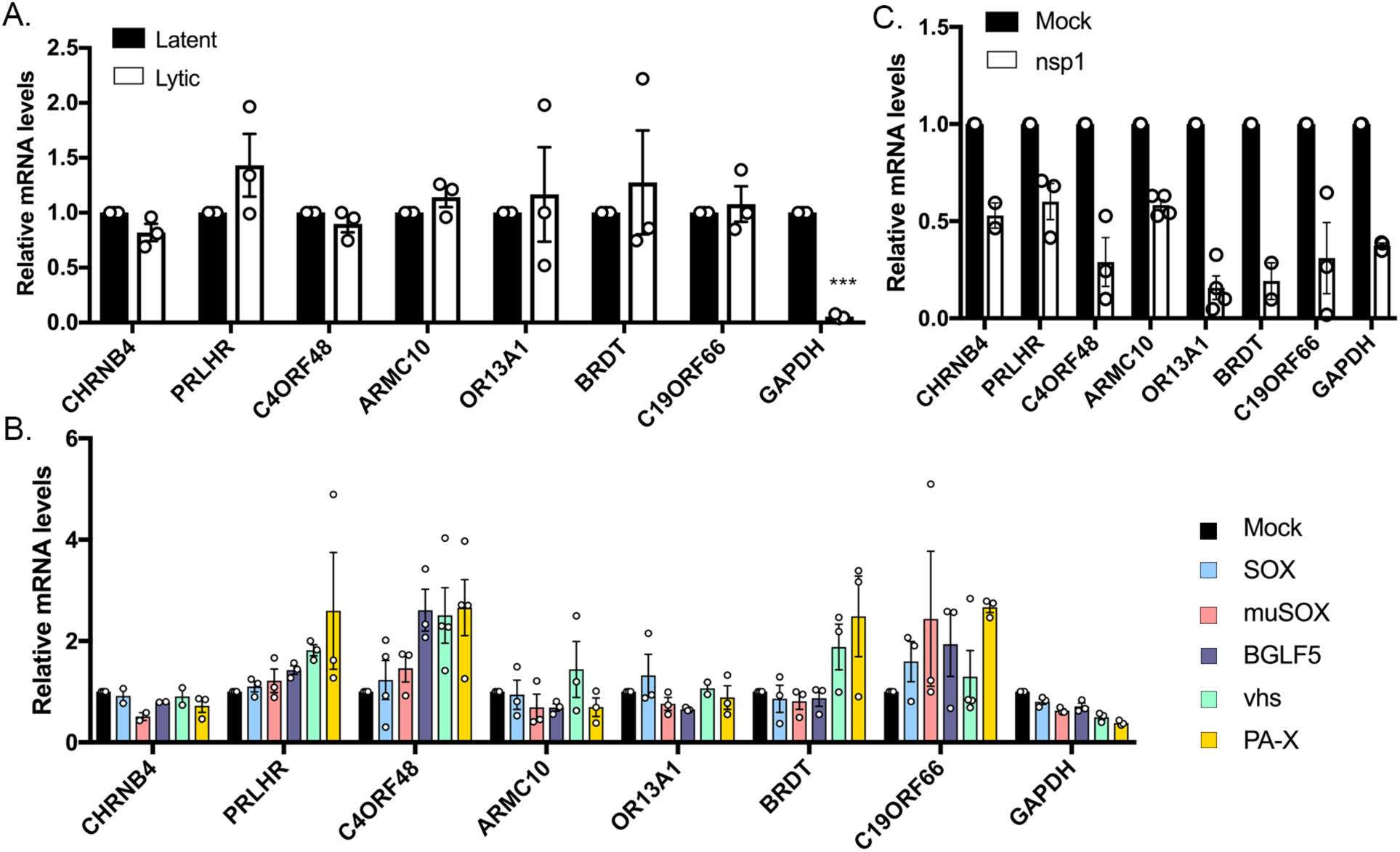
Top escapees identified by RNA-seq behave like SRE-containing transcripts. **(A)** Total RNA was extracted from unreactivated or reactivated KSHV-positive iSLK.219 cells and subjected to RT-qPCR to measure endogenous levels of the top candidates identified by RNA-seq. **(B)** 293T cells were transfected with an empty vector (Mock) or a plasmid expressing each of the viral endonucleases color coded on the right. After 24 h, total RNA was harvested and subjected to RT-qPCR to measure endogenous RNA levels. **(C)** 293T cells were transfected with an empty vector (Mock) or a plasmid expressing nsp1. After 24 h, total RNA was harvested and subjected to RT-qPCR to measure endogenous RNA levels.

### The C19ORF66 mRNA 3’ UTR contains an SRE

The pool of escaping transcripts did not appear to be strongly enriched for particular functions or processes when evaluated by GO-term analysis. We thus proceeded to manually mine the literature to identify functions that could be important during viral infection. We were drawn to C19ORF66 (also known as RyDEN, IRAV, and SVA-1), as it was reported to be an anti-viral interferon stimulated gene (ISG) in the context of multiple viral infections (40,43,44). Furthermore, the transcript for C19ORF66 appeared in our comparative RNA-seq as the top escapee in all the replicates and with all the endonucleases tested. We first evaluated whether this transcript contained a putative SRE-like element in its 3’UTR by testing whether it could protect the GFP mRNA, which is normally susceptible to viral endonuclease cleavage. We fused the C19ORF66 3’ UTR to GFP (C19-3’UTR) and found that it was sufficient to confer protection from SOX and other viral endonucleases in transfected 293T cells (**Fig. 3A**). Thus, similar to the IL-6 and GADD45B 3’ UTRs, previously identified dominant escapees, C19ORF66 contains an SRE-like element in its 3’UTR that is sufficient to provide protection against a range of viral endonucleases. As we previously demonstrated, there is no significant sequence conservation between the 3’UTRs of these known dominant escapees. However, the highest similarities were located near the second half of C19ORF66 3’UTR, and RNAfold secondary structure prediction of this UTR section revealed a long stem-loop structure with a bulge in the middle, consistent with previously found SRE structures (**Fig. S3**).

**Figure 3:**
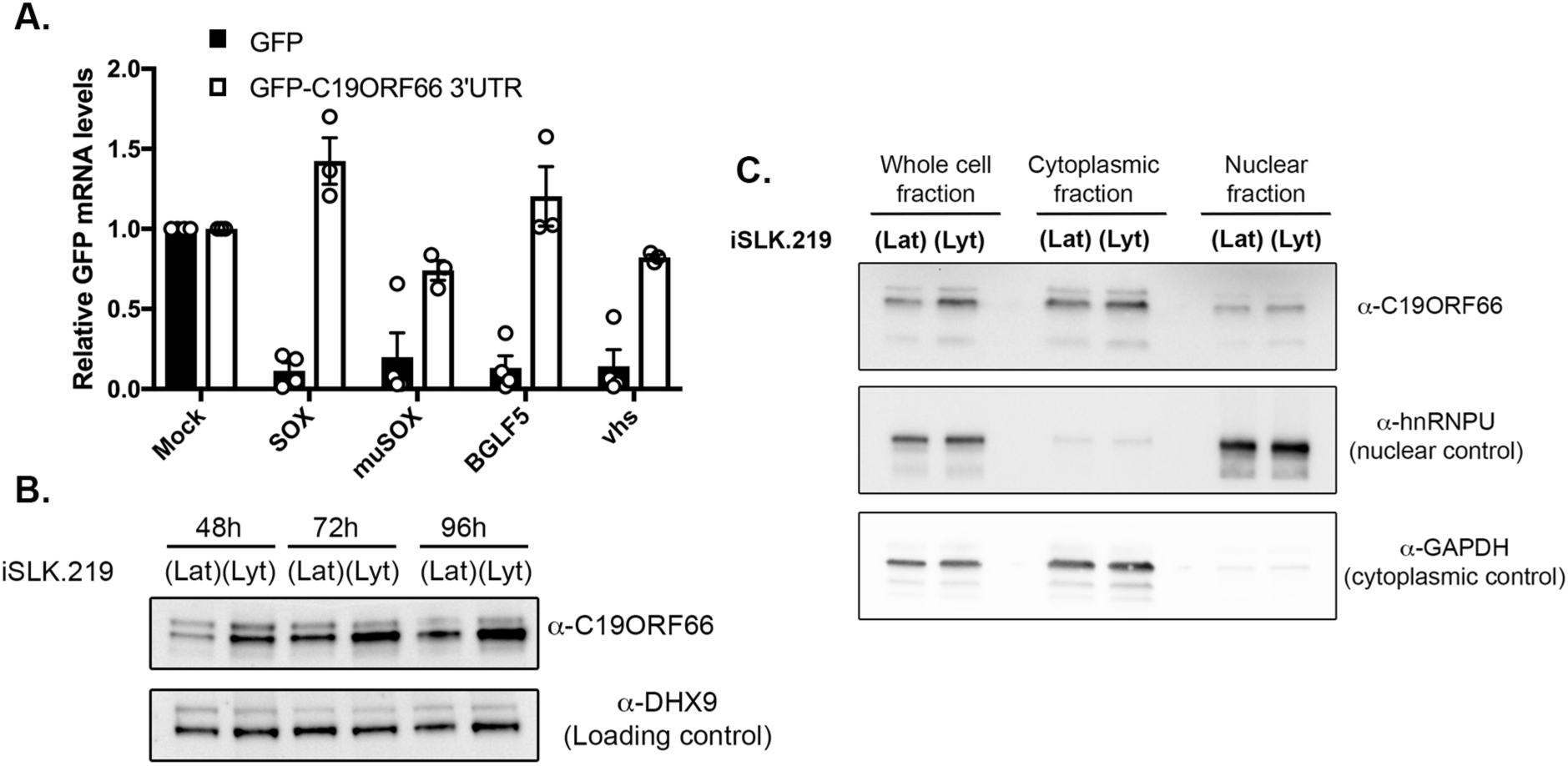
C19ORF66 mRNA is protected from herpesviral endonucleases by its 3’UTR and accumulates in the cytoplasm of iSLK.219 cells. **(A)** 293T cells were transfected with the indicated GFP reporter (GFP) or a GFP reporter containing C19ORF66 3’UTR sequence along with a control empty vector (mock) or a plasmid expressing SOX, muSOX, BGLF5 or vhs. After 24 h, total RNA was harvested and subjected to RT-qPCR to measure GFP mRNA levels. **(B)** KSHV-positive iSLK.219 cells were reactivated for the indicated times to induce KSHV lytic cycle (lyt) or not (KSHV latent phase maintained - lat). Cells were harvest, lyzed, resolved on SDS-PAGE and western blotted for the indicated antibodies. **(C)** Unreactivated (lat) or reactivated (lyt) KSHV-positive iSLK.219 cells were fractionated into nuclear and cytoplasmic fractions, and Western blotted for the indicated antibodies.

Because C19ORF66 expression was previously shown to be increased in the context of various viral infections, we next sought to investigate its expression upon KSHV lytic reactivation when host shutoff occurs. iSLK.219 cells were reactivated and total protein harvested at various time points over the course of 96 hours. C19ORF66 expression was increased upon KSHV lytic reactivation and continued to accumulate over time (**Fig. 3B**). Various other proteins were also previously shown to change subcellular localization in response to host shutoff (45), so we proceeded to monitor C19ORF66 expression and did not find differential shuttling upon KSHV lytic reactivation (**Fig. 3C**). Thus, C19ORF66 escapes SOX degradation by encoding an SRE-like element on its 3’ UTR that allows it to escape host shutoff and accumulate in lytically infected cells.

### C19ORF66 restricts KSHV infection

Given that C19ORF66 functions as an anti-viral protein during HIV and Dengue virus infection, we hypothesized that it could also play a role during KSHV infection. We thus further investigated the role of C19ORF66 in iSLK.219 cells. The recombinant KSHV.219 virus stably maintained in these cells constitutively expresses green fluorescent protein (GFP) from the EF-1 alpha promoter and can be used as a proxy for the presence of KSHV within cells. The KSHV.219 virus also encodes red fluorescent protein (RFP) under the control of the viral lytic PAN promoter (**Fig. 4A**). siRNA-mediated depletion of C19ORF66 in iSLK.219 cells during latency and at 48h and 72h post-reactivation was efficient, reducing expression levels by 94.6%, 97% and 97.8%, respectively (**Fig. 4B**). 72h hours post-reactivation, GFP and RFP positive cells were analyzed by fluorescence microscopy in siRNA C19ORF66-treated cells (or siRNA Control). C19ORF66 depletion resulted in a marked increase in the number of RFP positive cells (**Fig. 4C**). Conversely, overexpression of C19ORF66 in these cells (**Fig. 4D**) resulted in almost no RFP detection (**Fig. 4C**). Taken together, these results suggest that C19ORF66 expression negatively regulates the progression of KSHV life cycle.

**Figure 4:**
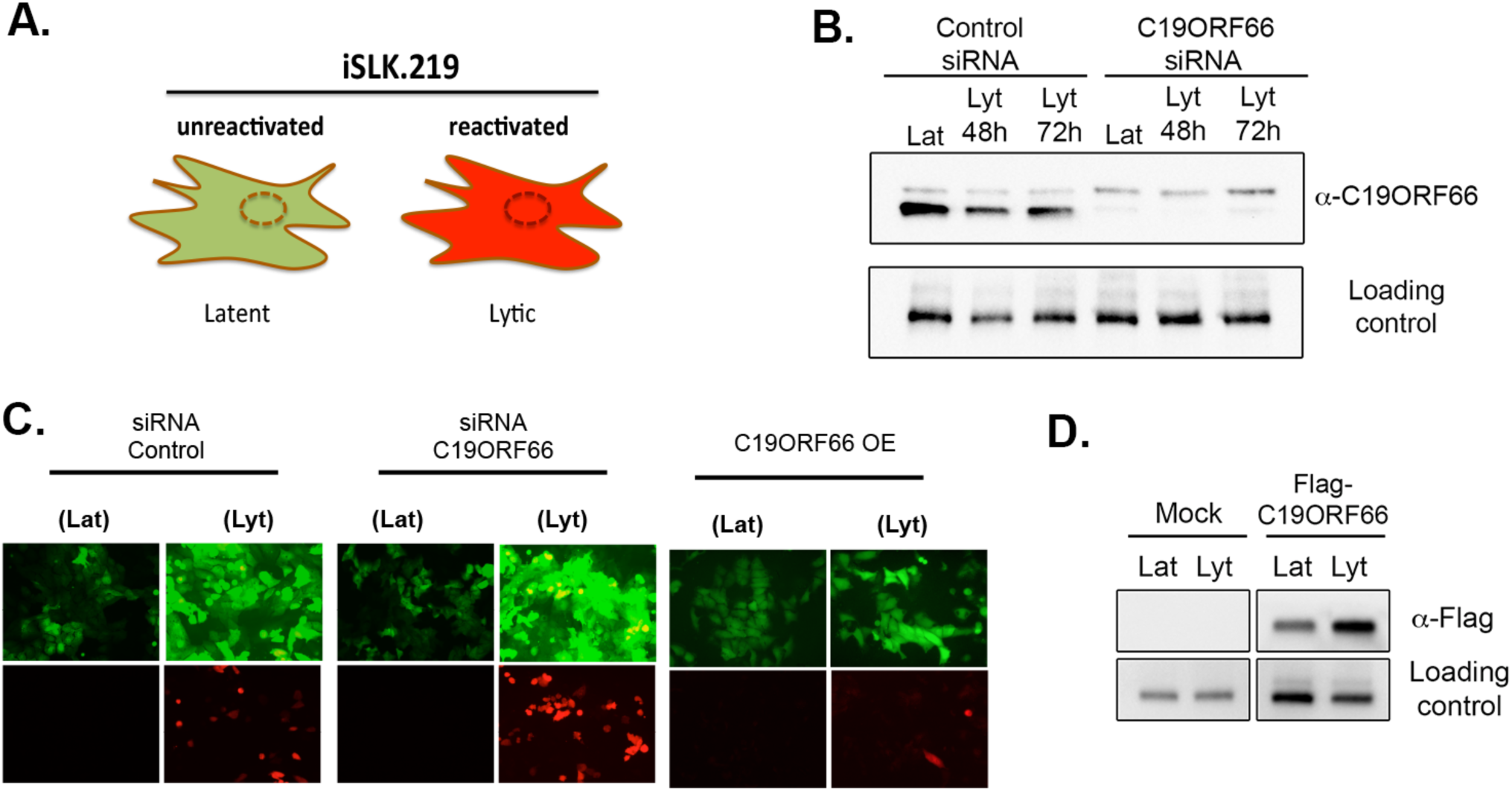
C19ORF66 restrict KSHV reactivation. **(A)** Diagram outlining the fluorescence pattern of iSLK.219 cells. **(B & D)** iSLK.219 cells were either treated with siRNAs targeting C19ORF66 (or control non-target siRNAs) for 48h (B) or transfected with a Flag tagged C19ORF66 (D). Cells were then reactivated with doxycycline and sodium butyrate, lyzed, and lysates were resolved on SDS-PAGE and western blotted with the indicated antibodies. (**C**) Cells treated with the indicated siRNA or transfected with C19ORF66 (Overexpression - OE) were checked for reactivation efficiency by monitoring the expression of GFP and RFP.

We next hypothesized that the reactivation defect due to C19ORF66 expression may lead to restriction of the formation of viral particles. To test this, we performed a supernatant transfer assay (**Fig. 5A**). iSLK.219 cells were treated with siRNA C19ORF66 (or control siRNA) and reactivated for 72h. Supernatants containing GFP expressing KSHV virions were collected and used to spinfect 293T cells (**Fig. 5B**). 24h later, we observed a higher number of GFP positive cells in the 293T cells infected with the supernatant coming from the iSLK.219 cells treated with the siRNA against C19ORF66. Since C19ORF66 seemed to affect important step in KSHV life cycle, we next assessed whether C19ORF66 also affects viral gene expression. Using RT-qPCR, we quantified the expression of several KSHV viral genes. Viral gene expression in KSHV unfolds as a cascade with the “early” (E) genes expressed right after lytic reactivation, followed by “delayed early” (DE) genes and finally, after viral replication, the “late” (L) genes. We harvested timepoints from 0 to 72h after iSLK.219 reactivation and measured RNA levels of genes representative of each gene class upon knock-down of C19ORF66. We observed a shift in viral gene expression with early and delayed early viral genes – but not with the late gene – which were expressed earlier and at higher levels in the C19ORF66 knocked down cells as measured by RT-qPCR (**Fig. 5C-E**). Taken together, these results suggest that C19ORF66 may restrict expression of certain early viral genes which in turn results in fewer newly formed viral particles being produced by KSHV infected cells.

**Figure 5:**
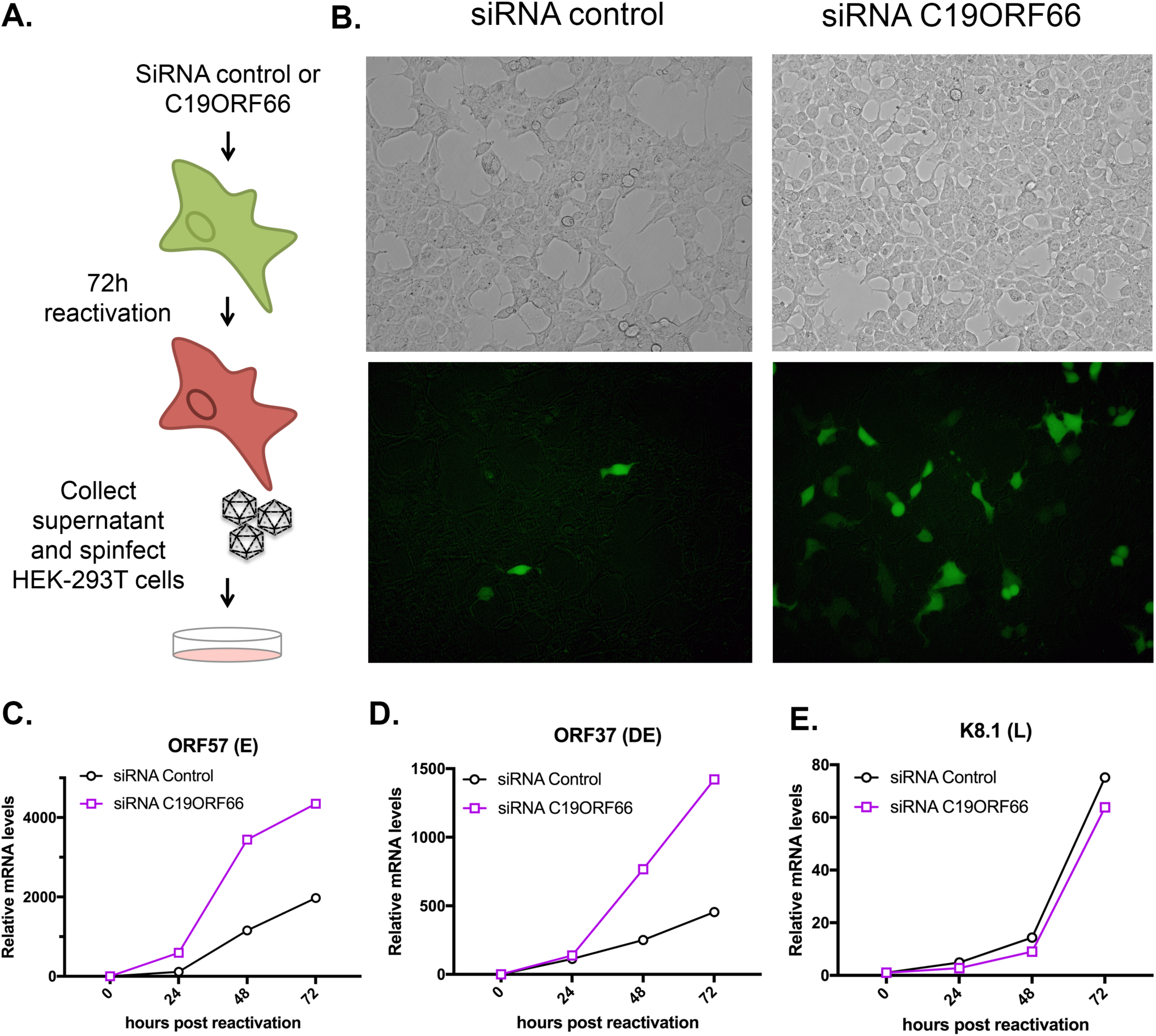
C19ORF66 knock down results in higher viral production yield and higher viral gene expression levels. (A) diagram depicting the supernatant transfer assay. (B) Supernatant transfer assay was used as a proxy for virion production and performed as described in A. Infection of 293T cells was monitored by imaging GFP on a fluorescent microscope. (C-E) Total RNA was extracted from iSLK.219 cells treated with siRNAs targeting C19ORF66 (or control non-target siRNAs) for 48h and reactivated for the indicated times. RNA was then subjected to RT-qPCR to quantify expression of the indicated viral genes.

## Discussion

Regulation of mRNA stability has emerged as a focal point for control of the host gene expression machinery. By accelerating RNA decay, viruses can increase their access to the host translation machinery and dampen the host response to infection. RNA degradation is often driven by virally encoded endonucleases that can target a wide array of mRNA by cleaving within a specific structured element (13,15). It is estimated that up to two thirds of total mRNA are degraded upon expression of these viral endonucleases (11,13,16). While recent studies have focused on *how* these viral endonucleases target mRNA, it remains unclear how and why some mRNA transcripts can escape viral-induced RNA decay. We previously demonstrated that certain transcripts escape by possessing in their 3’UTR an RNA element that protects them from viral endonucleases, while still allowing for normal RNA decay and cellular endonuclease cleavage (19-21). This raised a number of questions regarding how common these RNA escape elements are in the host genome and how their presence impacts the viral lifecycle. Here, we reveal that a cluster of 75 host transcripts can systematically escape viral-induced endonucleolytic cleavage. We hypothesize that these may contain similar RNA escape elements as the one we previously characterized in IL-6 and GADD45B and therefore could be important regulators of the viral-host interplay. IL-6 and GADD45B escape elements (referred to as SRE and G-SRE respectively) were shown to adopt a specific secondary structure that we hypothesized to be crucial in recruiting host protein to the 3’UTR of these escaping transcripts (19). This RNA-protein protective complex appears to be composed of core proteins as well as accessory proteins that may be transcript-dependent. One future goal is thus to expand our knowledge of the known escapees by exploring the RNA-protein complexes on these newly identified escaping transcripts with the hope of understanding the protein pre-requisite to forming a protective complex. More globally, determining whether such RNA elements impact RNA fate in uninfected cells will also be key in deciphering their role. To date, no such RNA elements have been found in viral genes, suggesting that this could be a cell specific mechanism that has evolved in response to viral infection.

No common functions were enriched in the pool of escaping transcripts, rendering it difficult to make any definite conclusion on whether these mRNAs escape degradation to benefit the host or the virus. Instead, we hypothesize that these spared mRNAs may have both pro and anti-viral functions. Furthermore, because of the large diversity of hosts infected by members of the herpesviridae, it would be interesting to investigate whether the orthologs of the escaping transcripts in other species also contain these RNA escape elements.

Here, we also characterized the top escaping transcript in our screen, C19ORF66. Through knock down and overexpression assays, our data indicate that C19ORF66 is restricting expression of KSHV early and delayed early genes, resulting in lower levels of viral reactivation and reduced yield of infectious viral particles. C19ORF66 is known to be upregulated in response to type I and type II IFNs (50,51) and to be upregulated in response to infection by a number of unrelated viruses (35-42). Furthermore, C19ORF66 was found to interact with the NS3 protein of Hepatitis C Virus (52), localize to the replication complex of DENV [33], and occasionally co-localize in the cytoplasmic compartment with HIV-1 Rev and Tat proteins (40), pointing to a potential conserved role for C19ORF66 as a key player in the host-pathogen response. While it is still unclear how C19ORF66 participates in the regulation of these viruses, it was hypothesized that it may be mediated through its interaction with PABPC and LARP, two major RNA binding proteins (43). PABPC and LARP were recently shown to be relocated upon SOX-induced widespread RNA decay and to be linked to the transcription feedback loop that occurs during host shutoff (45). PABPC in particular, was shown to be pivotal in triggering transcriptional repression in the nucleus after host shutoff, a process that favors expression of viral genes. It is therefore possible that C19ORF66, by interacting with PABPC, slows down PABPC relocalization to nucleus and restricts expression of viral genes. By Interacting with PABPC and LARP, C19ORF66 was also hypothesized to regulate decay of Dengue RNA by possibly influencing either translation or localization to p-bodies and stress granules (53) Determining whether C19ORF66 influences the PABPC shuttling pattern is an important future goal, as well as deciphering C19ORF66 interaction pattern upon KSHV infection and lytic reactivation.

Past literature on C19ORF66 has attributed C19ORF66 upregulation upon viral infection to interferon signaling. KSHV encodes multiple proteins that restrict the expression of interferon stimulated genes (ISG) (54) and yet, we observed an increased in C19ORF66 expression during KSHV lytic cycle. This suggests that in addition to escaping SOX-induced mRNA decay, C19ORF66 must have a mechanism to escape the KSHV encoded ISG inhibitors. This reinforces the idea that C19ORF66 is particularly important during KSHV infection and, to date, remains the only known ISG capable of escaping virally induced widespread mRNA decay.

Intriguingly, the viral endonucleases tested in this study come from both related and unrelated viruses, do not share the same targeting elements on their target mRNA, and are not known to be recruited to mRNA through similar pathways. It is thus notable that within the group of common escapees was one with a conserved anti-viral role. This underscores the utility of comparative approaches towards revealing broad regulators of viral infection.

Finally, none of the 3 known SREs (in IL-6, GADD45B, and now in C19ORF66) share significant sequence similarity, although they do all share similar predicted secondary structures. Thus, as predicted before, these RNA elements may function as scaffolds for recruiting a protective protein complex. Therefore, by manipulating the sequence of these RNA escape elements but maintaining the structure, these nuclease escape elements could be developed as tools to broadly inhibit viral endonucleases and open the possibility of turning these RNA elements into broad-acting anti-viral RNA therapeutics.

## Materials and Methods

### Cells and transfections

293T cells (ATCC) were grown in DMEM (Invitrogen) supplemented with 10% FBS. The KHSV-infected renal carcinoma cell line iSLK.219 bearing doxycycline-inducible RTA was grown in DMEM supplemented with 10% FBS (47). KSHV lytic reactivation of the iSLK.219 cells was induced by the addition of 0.2 μg/ml doxycycline (BD Biosciences) and 110 μg/ml sodium butyrate for 72 h.

For DNA transfections, cells were plated and transfected after 24h when 70% confluent using PolyJet (SignaGen). For small interfering RNA (siRNA) transfections, cells were reverse transfected in 6-well plates by INTERFERin (Polyplus-Transfection) with 10 μM of siRNAs. siRNAs were obtained from IDT as DsiRNA (siRNA C19ORF66: hs.Ri.C19orf66.13.1).

Fractionation experiments were performed following the REAP method (55). Briefly, cells were washed twice with ice-cold PBS and the cell pellet was lysed in 0.1% NP-40 PBS lysis buffer. The nuclei were then isolated by differential centrifugation at 10,000 x *g* for 10 sec and the supernatant retained as the cytoplasmic fraction. For western blotting, the nuclei were sonicated in 0.1% NP-40 PBS lysis buffer.

Supernatant transfers were carried in iSLK.219 cells. Cells treated with siRNA were reactivated with doxycycline and sodium butyrate for 72?h, supernatants were collected, filtered to remove any potential whole cells, and spinfected onto 293T cells at 1500rpm for 1 h at 37C. 24h later, cells were imaged on a fluorescent microscope.

### Plasmids

The C19ORF66 3’UTR was obtained as G-blocks from IDT and cloned into a pcDNA3.1 plasmid downstream of the GFP coding sequence. The C19ORF66 coding region was obtained as a G-block from IDT and cloned in a pcDNA4 Nter-3xFlag vector. All cloning step were performed using in-fusion cloning (Clonetech-takara) and were verified by sequencing.

### RT-qPCR

Total RNA was harvested using Trizol following the manufacture’s protocol. cDNAs were synthesized from 1 µg of total RNA using AMV reverse transcriptase (Promega), and used directly for quantitative PCR (qPCR) analysis with the SYBR green qPCR kit (Bio-Rad). Signals obtained by qPCR were normalized to 18S.

### Western Blotting

Cell lysates were prepared in lysis buffer (NaCl 150mM, Tris 50mM, NP40 0.5%, DTT 1mM and protease inhibitor tablets) and quantified by Bradford assay. Equivalent amounts of each sample were resolved by SDS-PAGE and western blotted with the following antibodies at 1:1000 in TBST (Tris-buffered saline, 0.1% Tween 20): rabbit anti-C19ORF66 (Abcam) rabbit anti-DHX9/RNA Helicase A (Abcam), rabbit anti-GAPDH (Abcam). Primary antibody incubations were followed by HRP-conjugated goat anti-mouse or goat anti-rabbit secondary antibodies (Southern Biotechnology, 1:5000).

### RNA-seq

Cells were transfected with constructs encoding fusion proteins between the herpesviral endonucleases (SOX, muSOX, BGLF5 and vhs) and the cell surface receptor Thy1.1 (CD90.1). Pure populations of cells expressing the endonucleases were obtained as describe before (45). Briefly, cells expressing the surface marker Thy1.1 were separated using the Miltenyi Biotec MACS cell separation system: transfected cells were incubated with anti-CD90.1 microbeads on ice for 15 min and magnetically separated according to the manufacturer’s instructions. RNA was then extracted from Thy1.1 positive cells by Trizol and purified as described above. Purity and integrity was assessed by bioanalyzer. After polyA selection, libraries were subjected to single-end sequencing on a HiSeq 4000. Read quality was assessed using fastqc. Using Galaxy (56), reads were then aligned to the human genome (hg38) by Bowtie2 and differential expression analysis were performed using Cufflink and Cuffdiff (57). For graphical representation in the heatmap, fold change values were saturated by an hyperbolic tan function with a cutoff set at 10. Hierarchical clustering was generated in Python using the SciPy package with complete linkage and Euclidian distance.

### Statistical analysis

All results are expressed as means ± S.E.M. of experiments independently repeated at least three times. Unpaired Student’s t test was used to evaluate the statistical difference between samples. Significance was evaluated with P values as follows: * p<0.05; ** p<0.01; *** p<0.001.

## Acknowledgments

We thank all members of the Muller Lab for their insights. We are grateful to the Glaunsinger lab for helpful discussions and to Ella Hartenian and Sarah Gilbertson for technical help with the Thy1.1 constructs. We would also like to thank Dr. Romain Vasseur for help with Python.

## Supporting Information

**Figure S1:** (**Left**) Volcano plot of all genes differentially expressed in Mock samples vs. Endonuclease expressing cells. Dots represent fold change and p-values as determined by CuffDiff. Significant fold change (-log10(p_value) of 0.001 and under) are highlighted in red. **(Right)** Distribution of fold change per endonuclease tested over mock sample and corresponding percentages on degrading transcripts.

**Figure S2:** Gene Ontology (GO) analyses performed using the Gene Ontology Consortium algorithm (http://www.geneontology.org) recapitulating the enriched functions found in the pool of escaping mRNAs per condition. The color scale represents the p-value of each GO term as assessed by the algorithm.

**Figure S3:** Sequence alignment and comparison of structure predictions obtained with RNAfold for C19ORF66 3’UTR with other known SRE transcripts: IL-6 and GADD45B.

**Table S1:** RNA-seq dataset. Summary table combining transcript ID and FPKM scores per condition (Mock sample, SOX, muSOX, BGLF5 and vhs).

**Table S2:** List of mRNA escaping all endonucleases tested by comparative RNA-seq as identified by hierarchical clustering. Each tab in this table represent the fold change over mock sample for each herpesviral endonuclease. Highlighted in yellow are the transcripts selected as top 10% for further investigation.

