## Supplementary figures and images for "C19ORF66 broadly escapes viral-induced endonuclease cleavage and restricts Kaposi’s Sarcoma Associated Herpesvirus (KSHV)"

### Figure S1

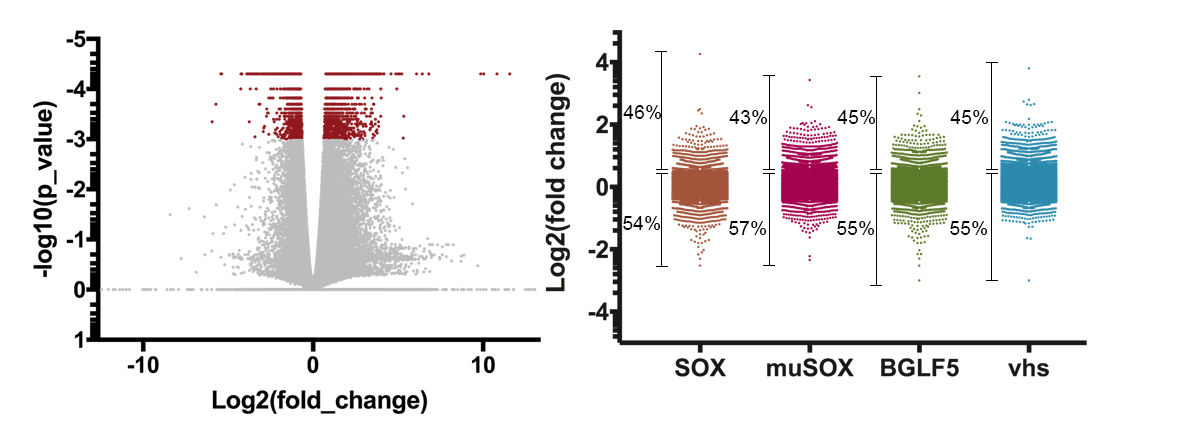

### Figure S2

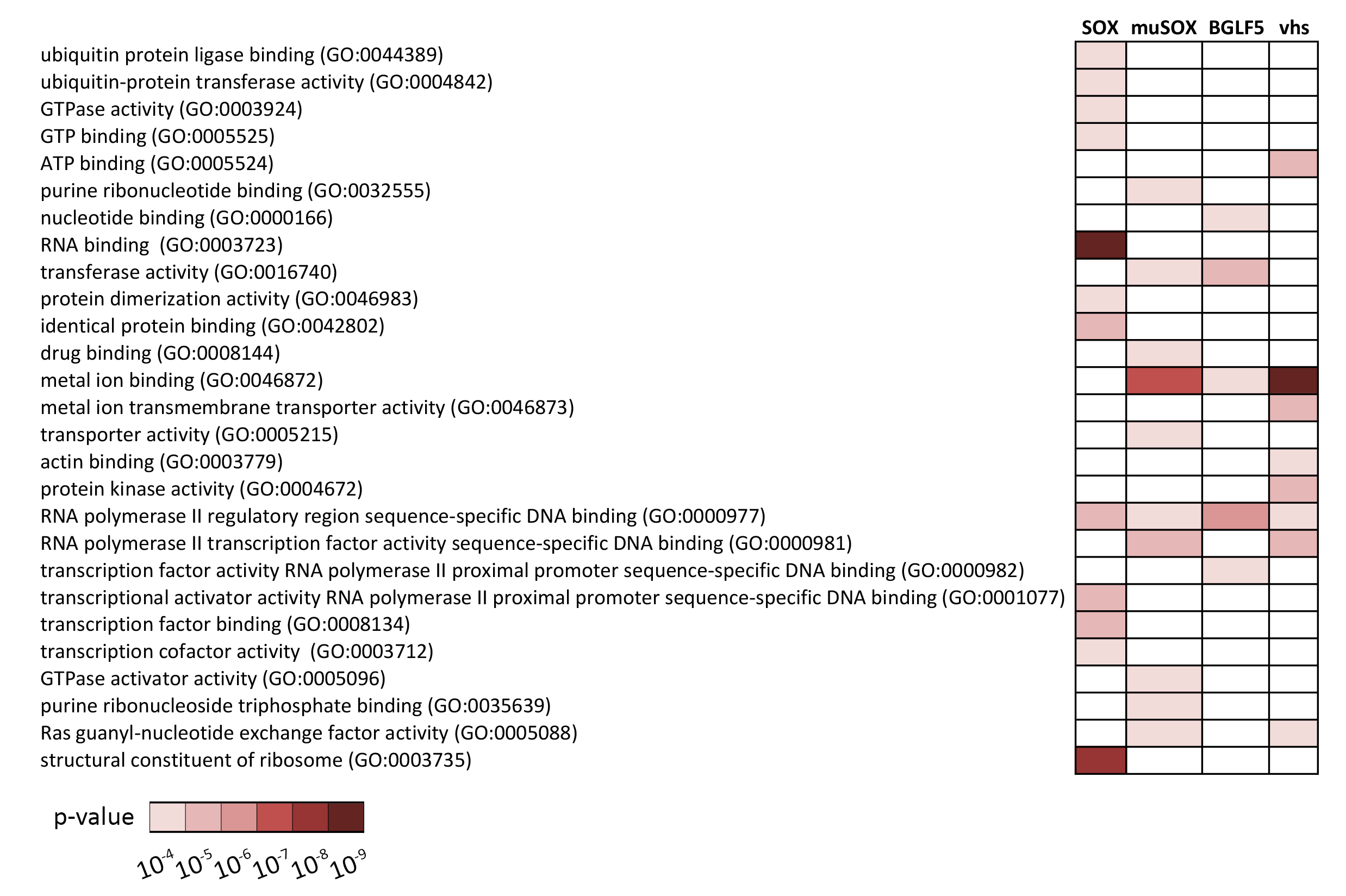

### Figure S3

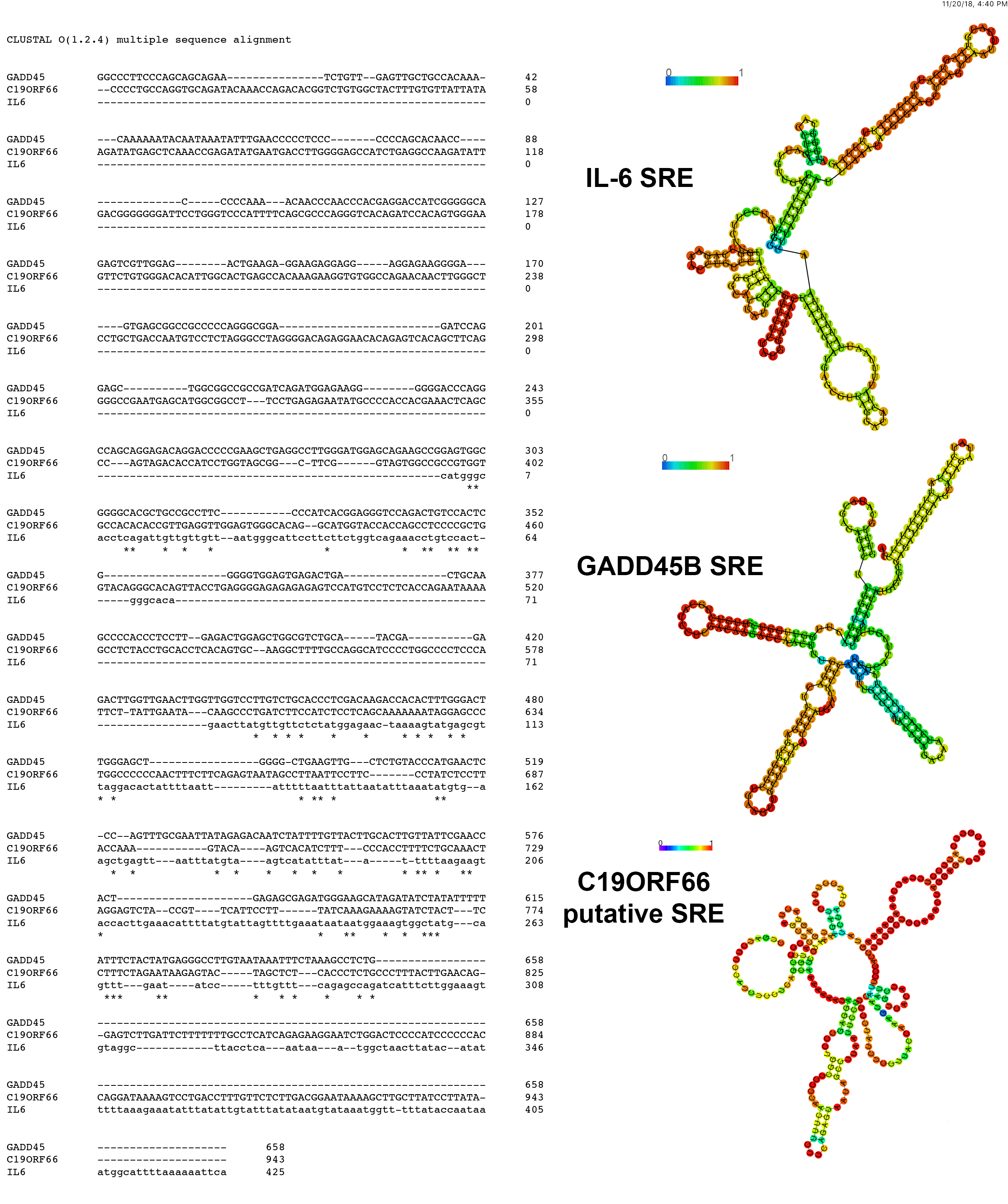
